# Sleep homeostasis reflects temporally integrated local cortical neuronal activity

**DOI:** 10.1101/756270

**Authors:** Christopher W. Thomas, Mathilde C. C. Guillaumin, Laura E. McKillop, Peter Achermann, Vladyslav V. Vyazovskiy

**Affiliations:** Department of Physiology, Anatomy and Genetics, University of Oxford, Oxford, UK; Nuffield Department of Clinical Neurosciences, University of Oxford, Oxford, UK; Institute of Pharmacology and Toxicology, University of Zurich, Zurich, Switzerland; The KEY Institute for Brain-Mind Research, Department of Psychiatry, Psychotherapy and Psychosomatics, University Hospital of Psychiatry, Zurich, Switzerland

## Abstract

The homeostatic regulation of sleep manifests as a relative constancy of its daily amount and intensity. Theoretical descriptions of this phenomenon define “Process S”, a variable with dynamics dependent only on sleep-wake history, whose levels are reflected in electroencephalogram (EEG) slow wave activity (0.5 – 4 Hz) during sleep. Here we developed novel mathematical models of Process S in mice, assuming that its dynamics are a function of the deviation of cortical neuronal firing rates from a locally defined set-point, crucially without explicit knowledge of sleep-wake state. Our results suggest that Process S tracks global sleep-wake history through an integration of local cortical neuronal activity levels over time. We posit that, instead of reflecting sleep-wake-dependent changes in specific variables and serving their homeostatic regulation, Process S may be a time-keeping mechanism which enables individuals to obtain a species-specific and ecologically-relevant quantity of sleep, even in the absence of external temporal information.

## Introduction

According to traditional theory, the need for sleep accumulates during wakefulness and dissipates during sleep. Despite decades of research, it is still uncertain precisely which biological variables form the substrate of sleep need, what characteristics of wake challenge their stability, how information about sleep-wake history is integrated over time, and how sleep mediates the restoration of homeostasis. Daily sleep amount varies greatly across the animal kingdom, yet individuals generally perform a relatively constant and species-specific amount of sleep each day. An even more fundamental question therefore remains whether homeostatic sleep regulation reflects an active process, dynamically shaping daily sleep architecture in response to a physiological need for the homeostatic regulation of specific variables, or whether it corresponds instead to an unknown innate time-keeping process which ensures only that a certain daily quota of sleep is obtained.

The earliest theories of sleep homeostasis supposed the existence of a single variable, termed Process S, which describes sleep drive at the global level (Borbély, 1982). This variable is assumed to always increase during wakefulness, independently of its content, and to decline during sleep. It is widely acknowledged that homeostatic sleep pressure is reflected in the levels of slow wave activity (SWA, 0.5-4 Hz spectral power) observable during NREM sleep in neurophysiological field potentials, such as electroencephalogram (EEG) or local field potential (LFP).

Current views on the origin of sleep homeostasis emphasise its local and activity-dependent component (Krueger & Tononi, 2011; Rattenborg et al., 2012; Tononi & Cirelli, 2014). It was shown that SWA is far from uniform across the brain, and that the behavioural and cognitive content of waking, beyond its mere duration, influences subsequent sleep and SWA (Kattler et al., 1994; Huber et al., 2004; Vyazovskiy & Tobler, 2008; Rector et al., 2009; Murphy et al. 2011; Fisher et al., 2016). Indeed, many candidate mechanisms for the substrate of sleep homeostasis implicate processes occurring at a cellular and network level. These include the maintenance of cellular homeostasis (Reimund 1994; Mackiewicz et al., 2007; Vyazovskiy & Harris, 2013; Bellesi et al., 2016), the replenishment of energy stores (Scharf et al., 2008), the influence of sleep-related signalling molecules such as adenosine or cytokines (Krueger et al., 2008), and the regulation of imbalanced synaptic strength (Tononi & Cirelli 2003; Vyazovskiy et al., 2008; Liu et al., 2010). Although the equivalence of these processes with Process S has not been conclusively demonstrated (Frank & Heller, 2019), the existing evidence supports the relevance of sleep-wake-dependent differences in neuronal activity for understanding the regulation of sleep.

Cortical neuronal activity is generally higher during waking compared to sleep (Vyazovskiy et al., 2009; Hengen et al., 2016; McKillop et al., 2018). Lower spike firing typical of sleep is due, at least in part, to regular periods of widespread synchronous network silence, termed off periods, intruding on ongoing activity (Steriade et al., 1993a; Steriade et al., 2001; Sanchez-Vives & Mattia, 2014). Importantly, off periods in neural populations are thought to underpin slow wave dynamics at the level of the field potential (Steriade et al., 1993b; Massimini et al., 2004; Buzsáki et al., 2012) and their properties reflect homeostatic sleep need (Vyazovskiy et al., 2009; McKillop et al., 2018; Saberi-Moghadam et al., 2018).

Neuronal firing rates typically fluctuate around a homeostatic set point, which is characteristic for individual cells and variable across the population (Turrigiano, 2011; Hengen et al., 2013; O’Leary et al., 2014; Styr et al., 2019). The homeostatic regulation of firing rates may depend on processes occurring specifically in sleep and wakefulness (Grosmark et al., 2012; Hengen et al, 2016) and evidence suggests that firing rates change as a function of time spent awake, conditional on the behaviour (Vyazovskiy et al., 2009; Fisher et al., 2016). Additionally, the magnitude and direction of state-dependent changes in firing rate differs between neurons, depending on the brain region and their individual firing rate set point (Miyawaki & Diba 2016; Watson et al., 2016; Miyawaki et al., 2019). Firing rate homeostasis may occur above the individual cell level (Slomowitz et al., 2015), perhaps through synaptic plasticity induced by the sleep slow oscillation which may act to narrow the population firing rate distribution (Levenstein et al., 2017).

Overall, the evidence suggests that the homeostatic regulation of sleep and of neuronal firing rates may be intrinsically related, however, this functional link remains incompletely defined. Mathematical modelling approaches present an opportunity to address this problem. To this end, we developed novel quantitative models of Process S and demonstrated that its temporal dynamics, on both a local and global level, can be derived entirely from local neuronal activity, without any reference to the animal’s sleep and wake states. We found that the magnitude of the deviation of multi-unit firing rate from a locally specified set point carries sufficient information to account for empirically derived patterns of SWA. We then introduced the total time spent in off periods as an alternative measure for Process S with more local origins than SWA, and showed how Process S dynamics may then be described in terms of two opponent processes, dependent on spiking rate and off period occurrence. Our data show that the global dynamics of Process S can be derived from a spatiotemporal integration of local neuronal activities. This suggests that a possible functional role for sleep homeostasis is to provide a precise intrinsic time-keeping mechanism, ensuring that individuals may obtain a species-specific daily quota of sleep even in the absence of external time cues.

## Materials & Methods

### Animals, surgery & husbandry

Chronic electrophysiological recordings from six male young adult (4.8 – 5.7 months old, mean 5.2 months) C57BL/6J mice were analysed here. This data set is a subset of that used in a previous study (McKillop et al., 2018). The animals were surgically implanted with electrodes for the continuous recording of electroencephalography, electromyography and cortical neuronal activity. EEG screw electrodes (Fine Science Tools) were inserted into the skull above the frontal cortex (primary motor area: anteroposterior 2mm, mediolateral 2mm) and occipital cortex (primary visual area: anteroposterior 3.5mm, mediolateral 2.5mm). Additional screw electrodes were placed contralateral to the occipital screw and above the cerebellum to serve as the ground and reference electrodes, respectively. A pair of stainless steel wires were inserted into the nuchal muscle for the recording of electromyogram (EMG). A polyimide-insulated tungsten microwire array (Tucker-Davis Technologies) was implanted through a craniotomy window into the frontal cortex (primary motor area: anteroposterior 2mm, mediolateral −2mm), contralateral to the EEG screw. The array comprised 16 wire channels of 33μm diameter, arranged in 2 rows of 8, with columnar separation of 250μm, row separation of 375μm and tip angle of 45 degrees. One row of wires was 250μm longer than the other to account for cortical curvature. A silicone gel (KwikSil, World Precision Instruments) was used to seal the craniotomy, and dental acrylic cement used to stabilise all the implanted electrodes. Surgeries were performed under isoflurane anaesthesia (4% induction, 1-2% maintenance). Analgesics were given immediately before surgery (1-2 mg/kg metacam and 0.08 mg/kg vetergesic, subcutaneous injection) and for at least 3 days during recovery following surgery (1-2 mg/kg metacam, oral). In addition, an immunosuppressant (0.2 mg/kg dexamethasone) was given the day before surgery, immediately before surgery and during recovery for at least 2 days. Animal wellbeing was closely monitored during recovery until a stable return to baseline was observed. All procedures were performed under a UK Home Office Project License and conformed to the Animal (Scientific Procedures) Act 1986.

Mice were housed individually following surgery. Two weeks after surgery, mice were transferred, still individually, to custom made Plexiglas cages (20.3×32×35 cm), containing a running wheel (Campden Instruments), which were placed within ventilated sound-attenuated Faraday chambers (Campden Instruments). The animals were exposed to a standard 12h-12h light dark cycle, with food and water available *ad libitum*. Mice were allowed to habituate to the recording chamber and to attachment of the recording cables for a minimum of three days.

### Experimental design

All the data analysed here were collected over two days. The first day served as a baseline, while the animals were completely undisturbed. A sleep deprivation protocol was enforced at light onset on the second day, immediately after the baseline day, and lasted 6 hours. Sleep deprivation was performed using the well-established gentle handling procedure (Fisher et al., 2016). During this period, experimenters constantly monitored both the behaviour and ongoing neurophysiological recordings of the mice. As soon as any animal showed signs of sleepiness (such as immobility, or slow waves in the EEG), novel objects were introduced to the cage (such as cardboard, colourful plastic and tissue paper) in order to encourage wakefulness. During the sleep deprivation period of 6 h, these mice slept 6.1 ± 2.9 min (mean ± sd) only.

### Data collection & pre-processing

Data acquisition was performed using a Multichannel Neurophysiology Recording System (Tucker Davis Technologies). EEG, EMG and microwire array LFP signals were filtered (0.1-100 Hz), amplified (PZ5 NeuroDigitizer preamplifier, TDT) and stored locally (256.9 Hz sampling rate). Custom written Matlab scripts were used for signal conversion and data pre-processing. The LFP, EMG and EEG signals were filtered again offline between 0.5-100 Hz (4^th^ order Type II Chebyshev filter) and resampled at 256 Hz.

Extracellular multi-unit spiking was additionally obtained from each microwire array channel, recorded at 25 kHz and filtered 300 Hz – 5 kHz. An amplitude threshold (at least 2 standard deviations, minimum −25 μV) was used to identify putative spikes. Individual spikes were saved as a voltage waveform comprising 46 data samples (0.48 ms before to 1.36 ms after threshold crossing) plus a time stamp of occurrence. Spiking activity from each channel was cleaned offline for artefacts using the Matlab spike sorting software Wave_clus (Quiroga et al., 2004). All putative single unit clusters identified by the algorithm from the same channel were merged, excluding only noise spikes. MUA firing rate for each channel was calculated in epochs of 4 s and expressed in Hz.

Vigilance states were scored manually by visual inspection at a resolution of 4 s (using the software SleepSign, Kissei Comtec). Vigilance states were classified as waking (low amplitude but high frequency EEG with high or phasic EMG activity), NREM sleep (presence of EEG slow waves, a signal of a high amplitude and low frequency, and a low level of EMG activity) or REM sleep (low amplitude, high frequency EEG, and low EMG). For this analysis, over the six mice, a total of 78 out of 96 channels could be analysed (minimum of 10 out of 16 from one mouse). The excluded channels were characterised by an unstable MUA firing rate, including a large drift in firing which was persistent and sleep-wake state independent. For the off occupancy models described below, 75 out of 96 channels were used; a further 3 channels had to be excluded because multi-unit firing rates, while stable, were too low to yield a reasonable estimate of the occurrence of off periods.

### Slow wave activity

Each EEG and LFP signal was processed to extract a measure of the slow wave activity (SWA). Signal segments were extracted within windows of 4-s duration and 1-s spacing (giving 3-s overlap), and Hann tapered. A Fourier transform was applied to each signal segment and the mean power in the slow wave range (frequencies from 0.5 to 4 Hz) was calculated. This measure was smoothed by finding the median over five temporally adjacent overlapping segments, yielding a SWA measurement for each sequential epoch of 4 s. For normalisation within each channel, SWA values were then expressed as a percentage of the mean SWA calculated over all artefact-free epochs scored as NREM sleep in the baseline 24 hours. The median value of the SWA during each continuous NREM sleep episode served as an estimate of Process S. NREM sleep episodes shorter than 1 min (15 epochs) were excluded, as in a previous study modelling Process S in mice (Guillaumin et al., 2018). Brief awakenings (short arousals accompanied by movement, lasting <20 s) were excluded from analysis but were not considered to be the ending of a NREM sleep episode.

### Off period detection and definition of “off occupancy”

One aim of this study was to develop measures of Process S independent of the EEG, and so we focused on off periods which are the neuronal counterpart of slow waves. Off periods refer to brief interruptions of spiking activity which occur synchronously across many recording sites, last approximately 70-100 ms (Vyazovskiy et al., 2009), and are coincident with the positive deflection of LFP slow waves (McKillop et al., 2018). In these studies, off periods were detected by pooling spikes over all channels and identifying inter-spike intervals (ISIs) that exceed a threshold duration. However, pooling spikes removes the ability to compare local differences, and if an off period does not involve all channels it will go undetected. To overcome this limitation, off periods were defined separately for each channel by looking at the co-occurrence of local slow waves and spiking silence.

There is no universally accepted method for slow wave detection, and a recent comparison suggests that a simple amplitude-threshold-based approach, while in theory adequate, may underperform due to channel differences in overall LFP amplitude and large amplitude fluctuations of higher frequencies (Bukhtiyarova et al., 2019). For this reason, the LFP was first filtered from 0.5 to 6 Hz (4^th^ order Butterworth filter), and a threshold defined individually for each channel, using the median plus one median absolute deviation of the peak amplitude of all positive half waves (including all vigilance states, but excluding epochs with artefacts). All positive half waves with peak amplitude above threshold were then considered to be slow waves.

Next, for each LFP slow wave, the multi-unit spike preceding and following the slow wave peak was identified and the corresponding ISI determined. The distribution of these ISIs which coincide with slow waves was often (in 64 out of 75 channels) unambiguously bimodal, allowing the threshold to be selected at the local minimum between these two modes. When there was no evidence of bimodality a value of 120 ms was chosen, corresponding to the maximum value selected for the other channels. All inter-spike intervals aligned to slow wave peaks with duration exceeding the threshold were considered off periods. Finally, the metric termed off occupancy was defined, for each epoch of 4 s, as the percentage of time spent in a detected off period during that epoch. Just as with SWA (see above), the median value of off occupancy over NREM sleep episodes was used to represent the level of Process S.

### Model fitting and parameter optimisation

Three theoretical models were used to describe the time course of Process S (see Results). The model equations were solved using a discrete time approximation, iteratively updating the value of modelled Process S in time steps of 4 s (Euler method). The fitting of various models to a particular channel of data is equivalent to finding the optimal choice of parameters to achieve the closest match between simulation and empirical SWA/off occupancy. In all models, the initial value of Process S was included as an additional free parameter. The selection of parameter values for model fitting was achieved using a semi-automated approach. First, an algorithmic methodology was established, which depends primarily on the definition of an error metric to assess fit quality between modelled Process S and empirical data. For each animal, NREM sleep episodes were identified with a duration of at least 1 minute. For each NREM sleep episode (***n=1:N***), the median empirical SWA/off occupancy (***X_n_***), and similarly the median modelled value of Process S (***S_n_***), were computed. The error metric (***E***) is defined as the sum of absolute differences between model and data, weighted by the relative episode duration (***w_n_***). This weight was defined as the absolute duration (***d_n_***) of the NREM sleep episode, normalised by the total duration of all episodes.

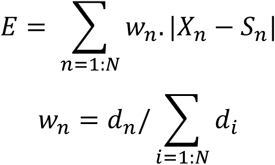

Algorithmic parameter optimisation was performed separately for each channel, aiming to minimise this error metric. This was achieved using the Matlab function *fminsearch*, which uses the Nelder-Mead simplex algorithm (Lagarias et al., 1998). Many parameter combinations produce very similar dynamics, and many possible optimal (or near-optimal) parameter regimes exist, therefore algorithmic optimisation is sensitive to initial values. For this reason, initial values for the parameters were first set manually, aided with the use of a custom-made Matlab graphical user interface. Manually selected values were then fed into the algorithmic optimisation. Final parameter values were visually inspected to ensure that this optimisation produced an improvement of fit.

The final fit quality of a model to the data was expressed as the median percent error (***E\****). This is calculated by finding the absolute difference between median empirical (***X_n_***) and simulated (***S_n_***) SWA/off occupancy for each NREM sleep episode, expressed as a percentage of the empirical SWA/off occupancy in that episode. ***E\**** is then defined as the median over all NREM sleep episodes of these difference values. This alternative error metric is used for presentation of results because it has more comprehensible units (percentage of empirical SWA/off occupancy) compared to the error metric, with arbitrary units.

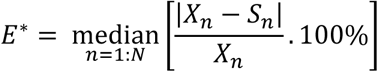

### Statistical analyses

The correlation between wake duration, wake firing rates and changes in SWA was calculated separately for each LFP channel. We first identified episodes of NREM sleep of at least 1-minute duration (exactly as for modelling) and obtained the median SWA in each episode. We then identified which intervening time periods, between two NREM episodes, comprised at least 80% wake and lasted at least 5 minutes. We then calculated Pearson correlation coefficients between i) the duration of these wake periods, ii) the mean multi-unit firing rate during these periods, and iii) the change in median SWA from preceding to the following NREM sleep episode.

Analysis of variance was performed to explore factors influencing model parameters and fit quality using the Matlab functions *anova1* (one-way) and *anovan* (two-way with unequal group size). For the rate parameters, ANOVA was calculated after applying a log transform. The effect size (***η^2^***) is calculated for a factor as its sum of squares divided by the total sum of squares in the ANOVA and reflects the fraction of the variance accounted for by that factor.

To summarise the results in figures, boxplots were included alongside individual data points. These indicate the median, 25^th^ and 75^th^ percentiles, with whiskers extending to the most extreme value which falls within 150 % of the interquartile range of the box. For these plots, results from individual channels were typically pooled across animals. In some cases, where indicated, channels from the same animal were presented as separate populations.

## Results

We selected a dataset of chronic recordings of frontal electroencephalogram (EEG) alongside local field potential (LFP) and multi-unit activity (MUA) spiking from primary motor cortex from six mice. These recordings were made continuously over 48 hours, starting at light onset, while the mice were freely behaving within their cage and exposed to a standard 12hr-12hr light-dark cycle. At light onset of the second day a sleep deprivation protocol was enforced for 6 hours, involving the presentation of novel objects. Outside of this sleep deprivation period, mice were undisturbed and able to sleep at will.

### Process S dynamics can be described as a function of vigilance state history

NREM sleep can be defined and distinguished from waking by the presence of high amplitude slow (0.5 - 4Hz) waves in the EEG. The average EEG spectral power in the slow wave range (termed slow wave activity; SWA) in NREM sleep varies as a function of the animal’s recent sleep-wake history, and this relationship has been captured in a classical quantitative theory using the concept of “Process S” (Borbély, 1982; Daan et al., 1984). Process S describes a variable whose magnitude can be estimated using the level of SWA during NREM sleep, reflecting the intensity of sleep, and which is interpreted as corresponding to the homeostatic component of sleep drive (Franken et al., 1991a; Achermann et al., 1993; Huber et al., 2000a; Vyazovskiy et al., 2007; Guillaumin et al., 2018). Theoretically, Process S follows simple dynamics: during wake it increases according to a saturating exponential function towards an upper asymptote, while during NREM sleep it decays exponentially towards a lower asymptote. There are many published variants of the precise equations for this model, but crucially, all these variants use the sleep-wake state history of the animal as the key predictive variable for Process S. Here, the specific equations used are:

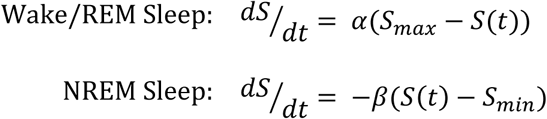

Where ***S*** represents the level of Process S, ***S_max_*** and ***S_min_*** are upper and lower asymptotes, and ***α*** and ***β*** are rate parameters. The first equation is applied when the animal is scored to be awake or in REM sleep and the second equation is applied when it is scored to be in NREM sleep. Here, we use the typical approach of applying the wake dynamics equation to REM sleep (Franken et al., 1991a; Huber et al., 2000a; Vyazovskiy et al.; 2007), but for simplicity do not set separate parameters for wake vs. REM sleep. For convenience, ***S*** is expressed in units equivalent to those of SWA. This classical formalism is later abbreviated as model Cl-SWA. Process S as described by these simple dynamics accounts for the time course of empirical SWA with high accuracy. We applied this model to SWA derived from the frontal EEG of all animals, and as expected obtained a high quality fit (Figure 1A).

**Figure 1.**
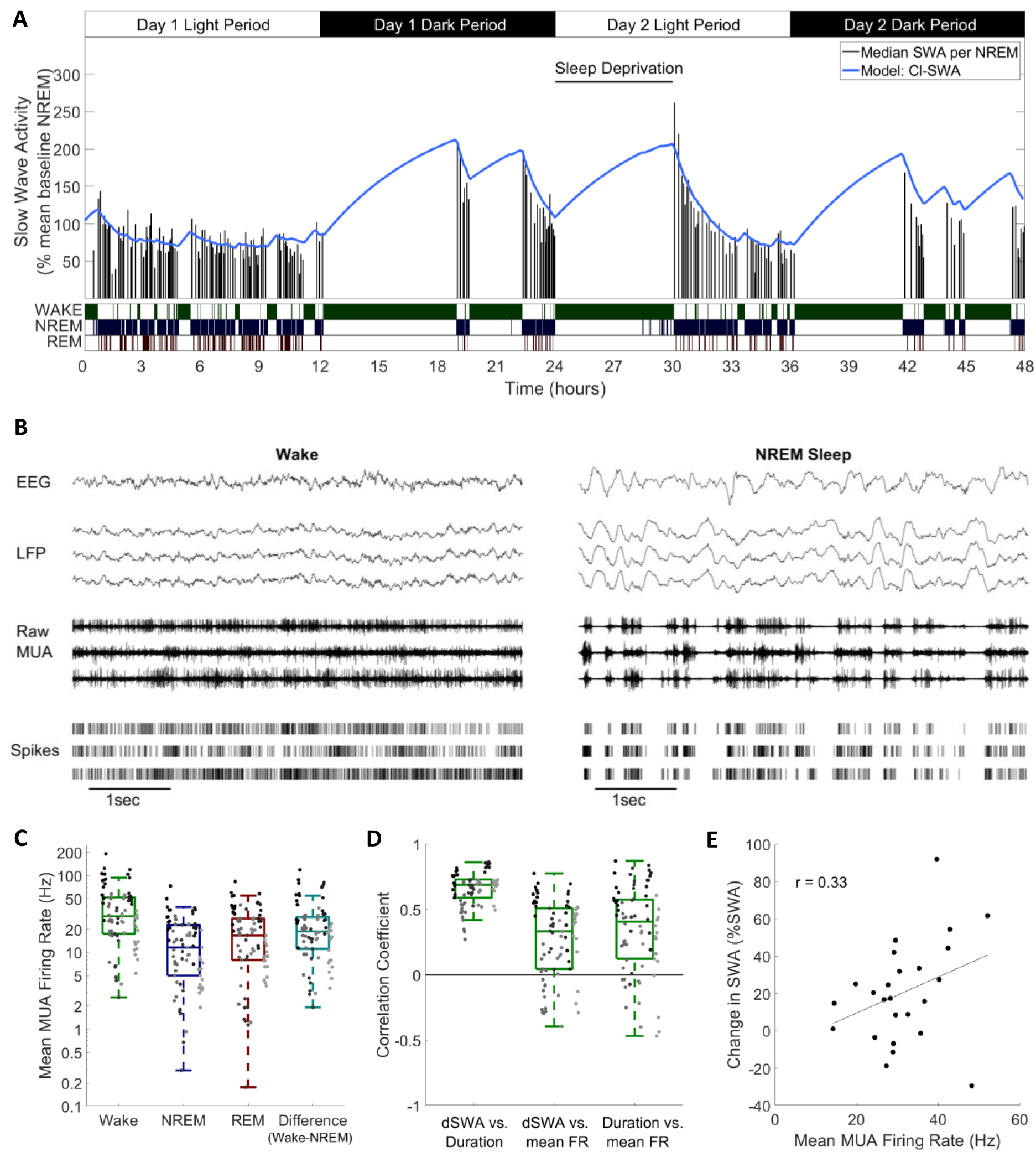
Cortical spike firing patterns are associated with the dynamics of Process S. **A)** An example of the classical state-based Process S model (blue) describing the dynamics of frontal EEG SWA (median per NREM sleep episode, black bars) over 48 hours in one representative animal. Sleep deprivation occurred as indicated at light onset of the second day and lasted 6 hours. Scored vigilance states are also shown. **B)** An example of frontal electroencephalogram (EEG), primary motor cortical local field potentials (LFP), corresponding raw signal with multi-unit activity (MUA), and detected spikes, in representative segments of waking and NREM sleep. Slow waves and synchronous spiking off periods are visible in NREM sleep but not in wakefulness. **C)** The distribution of mean firing rates during wake, NREM and REM sleep over the whole dataset, in addition to the difference in mean firing rates in wake compared to NREM sleep (all are positive, reflecting higher firing in wake). Points indicate channels grouped by animal (left to right), but boxplots reflect channels from all animals treated as one population. **D)** Distribution of correlation coefficients, calculated within each single channel, between wake episode duration (Duration), the change in slow wave activity (dSWA), and mean firing rate (mean FR). Points indicate channels grouped by animal (left to right), but boxplots reflect channels from all animals treated as one population. **E)** An example scatter plot of the correlation between the change in median SWA from one NREM episode to the next and the mean firing rate during the intervening period of wakefulness. This channel is representative because it has the median correlation coefficient of all channels.

### Neuronal firing rates are associated with Process S dynamics and vigilance state distributions

Whichever specific processes within the brain underpin these state-dependent dynamics for Process S, we hypothesised that it is associated in some way to neuronal spiking activity, which differs characteristically between wake and NREM sleep (Figure 1B). The occurrence of spiking off periods causes firing rates to be typically much lower during NREM sleep compared to wake and REM sleep. Consistent with this assumption, we found that the mean multi-unit firing rate averaged over all periods of wake was larger than the firing rate averaged over all periods of NREM sleep in every recording channel and every animal in this dataset (Figure 1C).

Because the slow oscillation is underpinned by local neuronal dynamics, it is expected to be highly heterogeneous across the neocortex. Regional differences in SWA dynamics have been previously described, for example between frontal and occipital EEG derivations (Werth et al., 1996; Huber et al., 2000b), and can be accounted for within the classical Process S model through the selection of locally variable values for model rate parameters (Zavada et al., 2009; Rusterholz & Achermann, 2011; Guillaumin et al., 2018). To explore whether spike firing rates might account for some of the variation in the rate of increase of Process S, we correlated the change in LFP SWA from one NREM sleep episode to the next, when separated by wakefulness lasting at least 5 minutes, with the mean spike firing rate during this intervening wake period, and with the duration of that wake period. Correlation coefficients were obtained separately for each recording channel, and pooled across animals. As expected, we obtained large positive correlation coefficients between the duration of a period of wakefulness and the change in LFP SWA in all channels (mean=0.66, sd=0.13; Figure 1D). Importantly however, the change in LFP SWA was on average also positively correlated with the firing rate (mean=0.27, sd=0.30; Figure 1D,E), meaning that, generally, a higher firing rate during waking was associated with a larger increase in SWA in subsequent NREM sleep. Interestingly, these phenomena were not independent, as a positive correlation was also found between wake episode duration and firing rate (mean=0.35, sd= 0.33; Figure 1D). These results further support the possibility that neuronal activity is associated with Process S dynamics. To test this hypothesis we next turned to a quantitative modelling approach.

### Process S dynamics can be described as a function of neuronal firing rates

We first sought to determine whether a model expressing Process S dynamics solely as a function of local multi-unit neuronal firing rates might describe the levels of sleep SWA in the corresponding LFP with comparable accuracy to the classical model which is dependent on vigilance states defined at the global level. The classical Process S model was used as a starting point for the development of a novel firing-rate-dependent alternative. To do this, the equations of the classical model were adapted in two ways. Firstly, an instantaneous firing rate threshold (***F_θ_***) was introduced as a new model parameter to replace the wake vs. sleep criterion, assuming an increase in Process S when the threshold is exceeded and decrease when firing is below. Conceptually, this firing rate threshold resembles a set point; a target firing rate at which the dynamics are stable. Secondly, we assumed that the rate of change of Process S is proportional to the difference between firing rate and this threshold. Introducing this change to the equations ensures that the rate of change of ***S*** is equal to zero exactly at the set point, and a continuous function of firing rate around this value. This version of the model is later abbreviated as Fr-SWA. The equations are:

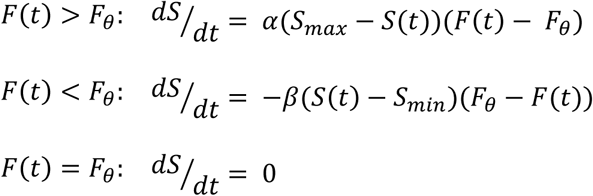

We applied this novel model, alongside the classical model, to describe SWA dynamics at the LFP level, by finding parameter values that would minimise the difference between empirical SWA and modelled Process S. The two models were fit to the SWA from each LFP channel separately, using the multi-unit firing rate from the same channel for the firing-rate-dependent model. The models were also fit to the SWA obtained by averaging the LFP (and firing rate) over the whole population of channels within the same mouse.

Figure 2 shows two examples of the fit of both models, first to SWA derived from the average LFP, and also to SWA derived from a single LFP channel (these examples are from the same animal as in Figure 1A). The overall pattern of Process S dynamics was similar at both LFP and EEG recording levels and was well described by both the classic and novel firing-rate-based model. Throughout the dataset, this purely firing-rate-dependent model described the overall dynamics of LFP SWA during NREM sleep to a comparable accuracy as the classical model, as reflected in the median percent error deviation between modelled and empirical SWA (Figure 3B). Both the model type and animal have a highly significant effect on the median percent error of the model fit to individual LFP channels, although there is no significant interaction (Model: F_(1, 144)_ = 25.9, p = 1.1×10^−6^; Animal: F_(5, 144)_ = 30.2, p = 6.3×10^−21^; Model × Animal: F_(5, 144)_ = 1.65, p = 0.15; two-way ANOVA unequal groups). Errors were higher for the novel model, importantly however, the differences in fit quality due to the model type are small relative to the effect size of the particular animal and channel, on which fit quality depends much more strongly (Model: η^2^ = 0.079; Animal: η^2^ = 0.459; Model x Animal: η^2^ = 0.025; Channel (residuals): η^2^ = 0.437). The errors of the model fit on the averaged LFP and on the EEG (classical model only) are also shown in Figure 3B, and are more explicitly compared in Figure 3C. We do not find any significant effect of the field potential level (EEG, average LFP, single LFP channels) on the model error (F_(2,171)_ = 0.2, p = 0.82, one-way ANOVA).

**Figure 2.**
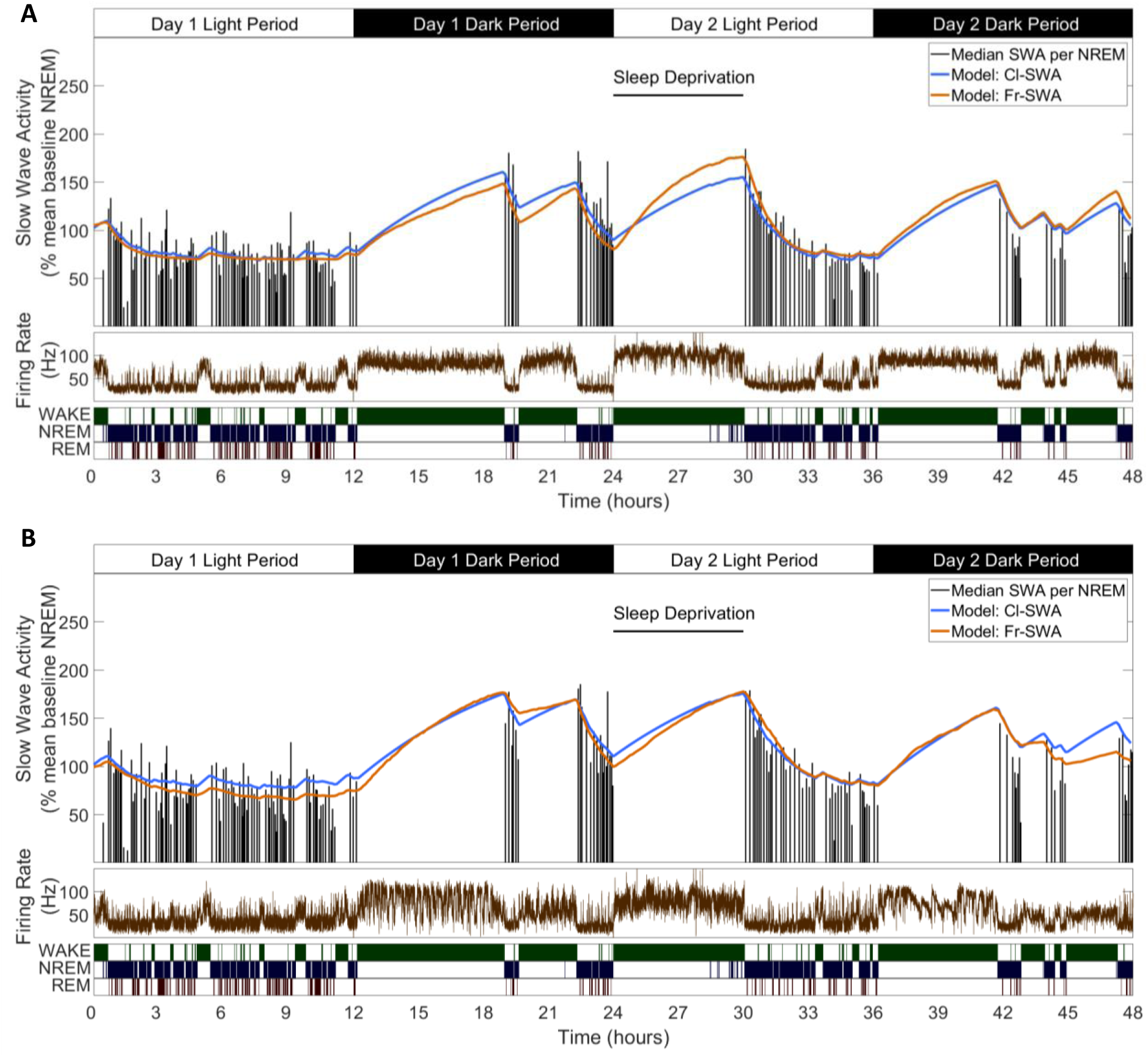
Slow wave activity dynamics at the LFP level can be modelled using multi-unit spiking information. **A)** An example from one representative animal modelling the SWA averaged over all LFP channels, of both the classical model (blue) and novel firing-rate-based model (orange), calculated from the firing rate also averaged over all LFP channels (brown). **B)** An example of both models applied to the SWA of a single LFP channel, which came from the same animal as used in **A**.

**Figure 3.**
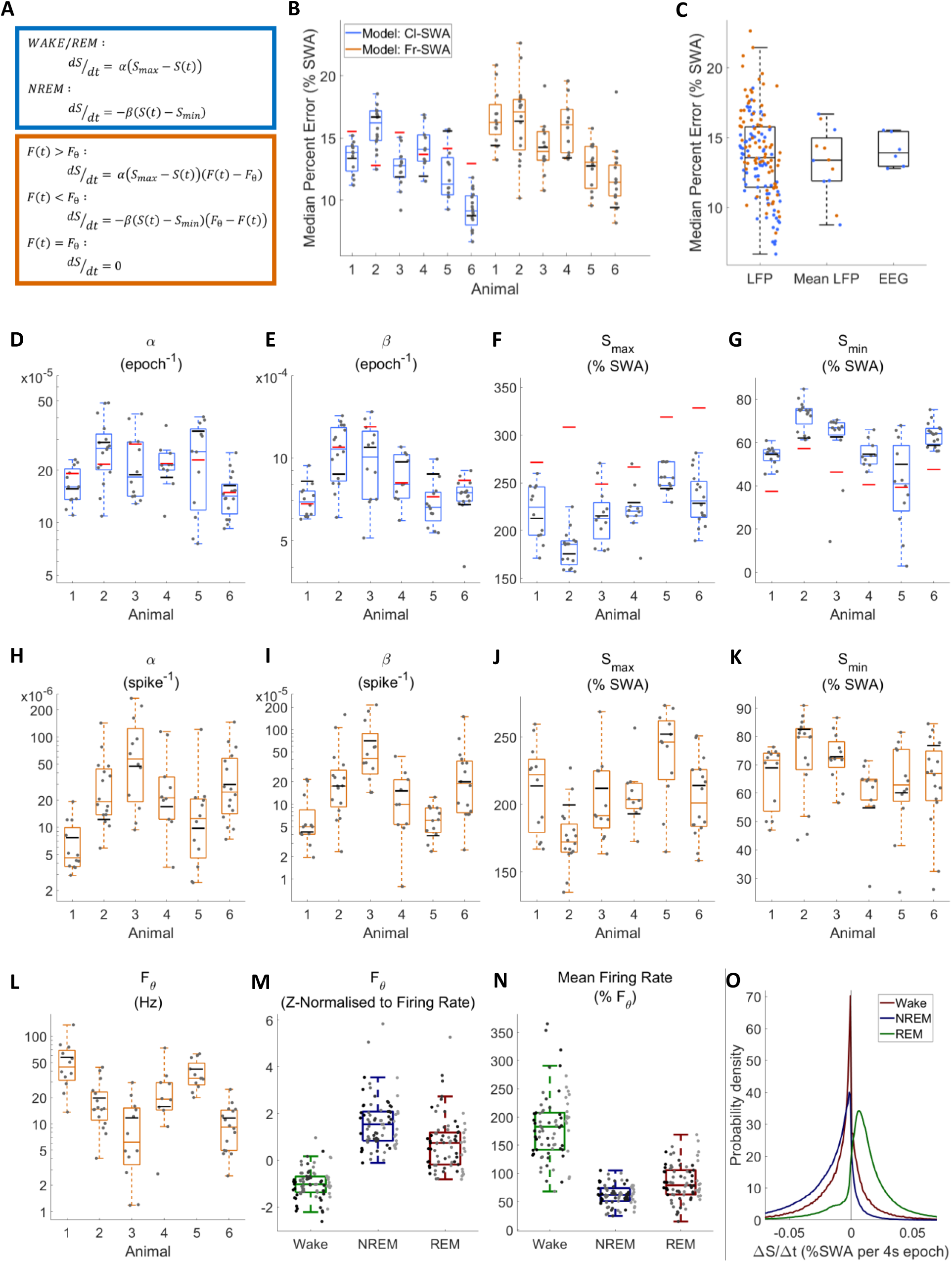
The fit quality and parameters for both classical and firing-rate-based models of LFP SWA. **A)** Equations for the classic state-based model (blue) and novel firing-rate-based model (orange). **B)** For each animal, the distribution over channels of the median difference between the model and empirical SWA, expressed as a percentage of empirical SWA, for both classic and firing-rate-based models. **C)** The same median percent error, grouped over animals and models, separately showing errors at the level of single LFP, averaged LFP and EEG. **D-G)** The distribution of values used for each parameter in the classic model and **H-L)** in the firing-rate-based model, with boxplots plotted separately for each animal. The black lines indicate the values obtained from modelling the averaged LFP SWA and firing rate over all channels within an animal. Additionally, the red lines in **D-G** give the parameter value obtained from the model of the frontal EEG SWA of that animal. **M)** The distribution of the final optimised value for the firing rate set point parameter (***F_θ_***) of the firing-rate-based model, z-normalised to the distribution of firing rates within wake, NREM and REM sleep. Points indicate channels grouped by animal (left to right), but boxplots reflect all channels treated as one population. **N)** The distribution of mean firing rate in wake, NREM and REM sleep, expressed as a percentage of the firing rate set point parameter (***F_θ_***). Points indicate channels grouped and coloured by animal, but boxplots reflect all channels treated as one population. **O)** The distribution of the change in Process S (***ΔS/Δt***) from one 4-s time step to the next derived from the Fr-SWA model in wake, NREM sleep and REM sleep. All mice, channels and time are pooled. Source data is provided for this figure which contains all simulated Process S time courses, model parameters and the data required to reproduce them.

The distributions of final optimised parameters are shown in Figure 3D-G for the classic model and Figure 3H-L for the firing-rate-based model. Most parameters in both models were significantly different between animals, with the exception of ***S_min_*** in the firing-rate-dependent model (Cl-SWA ***α***: F_(5,77)_ = 4.03, p = 2.8×10^−3^; Cl-SWA ***β***: F_(5,77)_ = 8.65, p = 1.9×10^−6^; Cl-SWA ***S_max_***: F_(5,77)_ = 14.35, p = 9.8×10^−10^; Cl-SWA ***S_min_***: F_(5,77)_ = 12.42, p = 1.1×10^−8^; Fr-SWA ***α***: F_(5,77)_ = 8.33, p = 3.0×10^−6^; Fr-SWA ***β***: F_(5,77)_ = 9.27, p = 7.5×10^−7^; Fr-SWA ***F_θ_***: F_(5,77)_ = 14.3, p = 1.0×10^−9^; Fr-SWA ***S_max_***: F_(5,77)_ = 6.64, p = 3.9×10^−5^; Fr-SWA ***S_min_***: F_(5,77)_ = 2.16, p = 0.07; one way ANOVA). The optimised rate parameters for EEG and averaged LFP data consistently fall within the range of the single channel population (Figure 3D-L). The relationship between the optimal firing set point and state specific firing rates is shown in Figure 3M. Firing rate threshold was z-normalised (subtract the mean firing rate over a given state and divide by the standard deviation) separately with respect to the distribution of firing rates in wake, NREM and REM sleep. This shows, as expected, that the set point is typically below mean firing in wake (−1.0 ± 0.6; mean ± sd), well above mean firing in NREM sleep (1.6 ± 1.1), and slightly above mean firing of REM sleep (0.7 ± 1.1). These were all significantly different from zero (wake: p = 1.6×10^−13^, NREM: p = 1.9×10^−14^, REM: p = 3.4×10^−7^, two-sided Wilcoxon signed rank test). Figure 3N further shows the state-dependent distribution of average firing rate, expressed as a percentage of the chosen firing set point parameter. Mean firing was 182 ± 57.6% of the firing rate set point during wake, 61.8 ± 17.6 % during NREM sleep, and 84.2 ± 31.8 % during REM sleep. Again, these distributions were all significantly different from 100% (Wake: p = 6.7×10^−14^, NREM: p = 2.0×10^−14^, REM: p = 5.4×10^−5^). Note that, in a few channels, the optimal firing rate threshold is actually above the mean firing rate during waking. This occurs when the waking firing rate distribution overlaps substantially with the NREM sleep firing rate distribution but has a heavier tail. REM sleep, in this dataset, is typically associated with firing below the set point, and therefore Process S decreases, albeit at a reduced rate compared to NREM sleep (Figure 3O). Interestingly, there are no significant correlations between the model fit error and the threshold normalisation relative to any one vigilance state (Wake: p=0.06, r=0.21; NREM: p=0.35, r=-0.11; REM: p=0.94, r=-0.01), suggesting that there is no clear relationship between the firing rate set point and firing rate distribution of any one particular state.

### Process S can be defined in terms of neuronal spiking off periods

When considering the relationship between multi-unit firing rates and LFP slow waves, a conceptual complication arises due to the different origins of the LFP and MUA from within the same channel. While the MUA firing rate represents the activity of only a few individual neurones, factors such as volume conduction result in spatial smoothing of the LFP signal, and as such it represents the combined activity of neurones covering a cortical area potentially on the order of several millimetres (Kajikawa & Schroeder, 2011). This means that when a slow wave is detected in the LFP, it is not guaranteed that all local neurones contributing to the MUA are necessarily silent (Todorova & Zugaro, 2019). Similarly, not every long interspike interval occurs during a slow wave. An estimation of the occurrence of off periods may be obtained by combining LFP and spiking data (Figure 4A). Slow wave detection was performed on each LFP over the whole 48 h, including all vigilance states (0.5-6 Hz filter followed by an amplitude threshold, values shown in Figure 4C). All multi-unit inter-spike intervals (ISIs) which coincide with the peak of detected slow waves were identified. The distribution of the duration of these ISIs was often (64 out of 75 channels) unambiguously bimodal (Figure 4B). This could be interpreted as evidence of the existence of two spiking states occurring locally during the more widespread network slow oscillation; high frequency spiking (on period) and extended silence (off period). Note that these two distributions do not simply correspond exactly to sleep vs. wake conditions, because bimodality is often evident in the distribution of slow wave coincident ISIs from NREM sleep only, or even from REM sleep only (Figure 4B). The distribution of slow wave coincident ISIs (over all vigilance states) was used to define an ISI duration threshold for the detection of off periods, separately for each channel. The values used for this threshold are shown in Figure 4D. The average multi-unit firing rate aligned to the peak of slow waves in detected off periods reveals a clear suppression of firing, consistent with expectations (Figure 4E). A rebound increase in average firing is visible in this example channel immediately after the off period, as has been previously documented in some cortical neuronal populations (Chauvette et al., 2010). The total fraction of time each channel spends in off periods was calculated over all epochs of 4 s, and termed the “off occupancy”. Off occupancy defined in this way is high in NREM sleep and low in both wake and REM sleep (Figure 4F). The existence of a non-zero frequency of local cortical off periods has been previously reported during wakefulness (Vyazovskiy et al., 2011; Vyazovskiy et al., 2014; Fisher et al., 2016), and REM sleep (Funk et al., 2016). The off occupancy measure displays similar temporal dynamics to LFP (and EEG) SWA over both sleep deprivation and spontaneous sleep and wake (Figure 4G).

**Figure 4.**
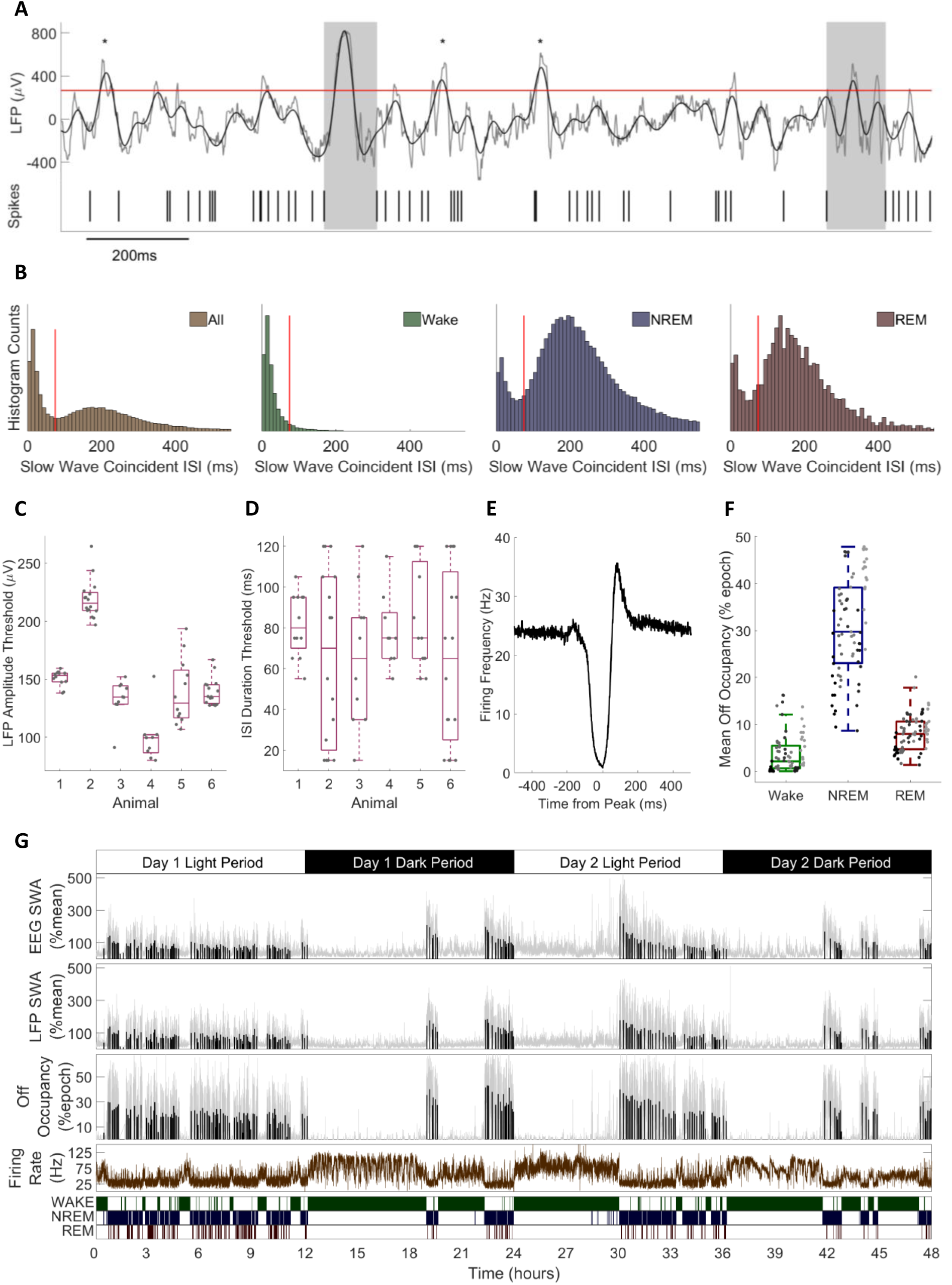
The definition of off periods and off occupancy. **A)** An example section of LFP (raw in grey, 0.5-6Hz filtered in black) and simultaneous MUA spikes. During this time window, the filtered LFP crosses above the amplitude threshold (265 μV for this channel, red line) five times. The multi-unit inter-spike interval aligned to two of these peaks exceeds the duration threshold (85ms for this channel) and so two off periods are detected (grey boxes). ISIs aligned to the other three out of five crossings (asterisks) are too short to be considered off periods. **B)** Histograms of multi-unit inter-spike intervals aligned with detected slow waves for this example channel. The four plots show, from left to right, ISIs over the whole recording and ISIs separately during wake, NREM and REM sleep only. The ISI duration threshold (red line) is selected using the histogram of all ISIs (leftmost) at the minimum between the two modes. **C)** The distribution of LFP amplitude and **D)** ISI duration threshold values used for definition of off periods for each channel, with boxplots plotted separately for each animal. **E)** The mean multi-unit firing rate over a period of 1 s centred on the peak of detected slow waves, calculated over all slow waves within one example channel with a resolution of 1 ms. **F)** Distributions of mean off occupancy (%) for all channels averaged over wake, NREM and REM sleep. Points indicate channels grouped by animal (left to right), but boxplots reflect all channels treated as one population. **G)** Off occupancy is shown alongside EEG and LFP SWA for an example channel over 48 hours. Traces represent these values calculated at 4-s resolution (light grey), in addition to the median value per NREM sleep episode, as used for model fitting (black bars). Firing rate (brown) and scored vigilance states are also shown.

### Process S dynamics, defined using off periods, can be described as a function of vigilance states or neuronal firing rates

In order to investigate whether the off occupancy measure reflects Process S, we applied the classical state-based model to the time course of off occupancy, exactly as was done with single channel SWA. The classical model was applied with its equations unchanged, and is abbreviated as Cl-Off. Figure 5A shows an example of the resulting Process S time course obtained in this way with a high quality of fit. In contrast, some changes were introduced to the firing-rate-based model in order to describe off occupancy dynamics. We considered that the different dynamics above vs. below a particular firing rate set point may be due to two opponent processes simultaneously active in dynamic opposition but with differential magnitude in wake vs. NREM sleep. Specifically, we consider a Process S increasing term which is proportional to instantaneous firing rate, and a Process S decreasing term which is proportional to the time spent in off periods (off occupancy). The equation used is:

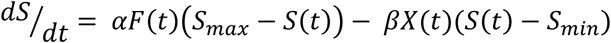

**Figure 5.**
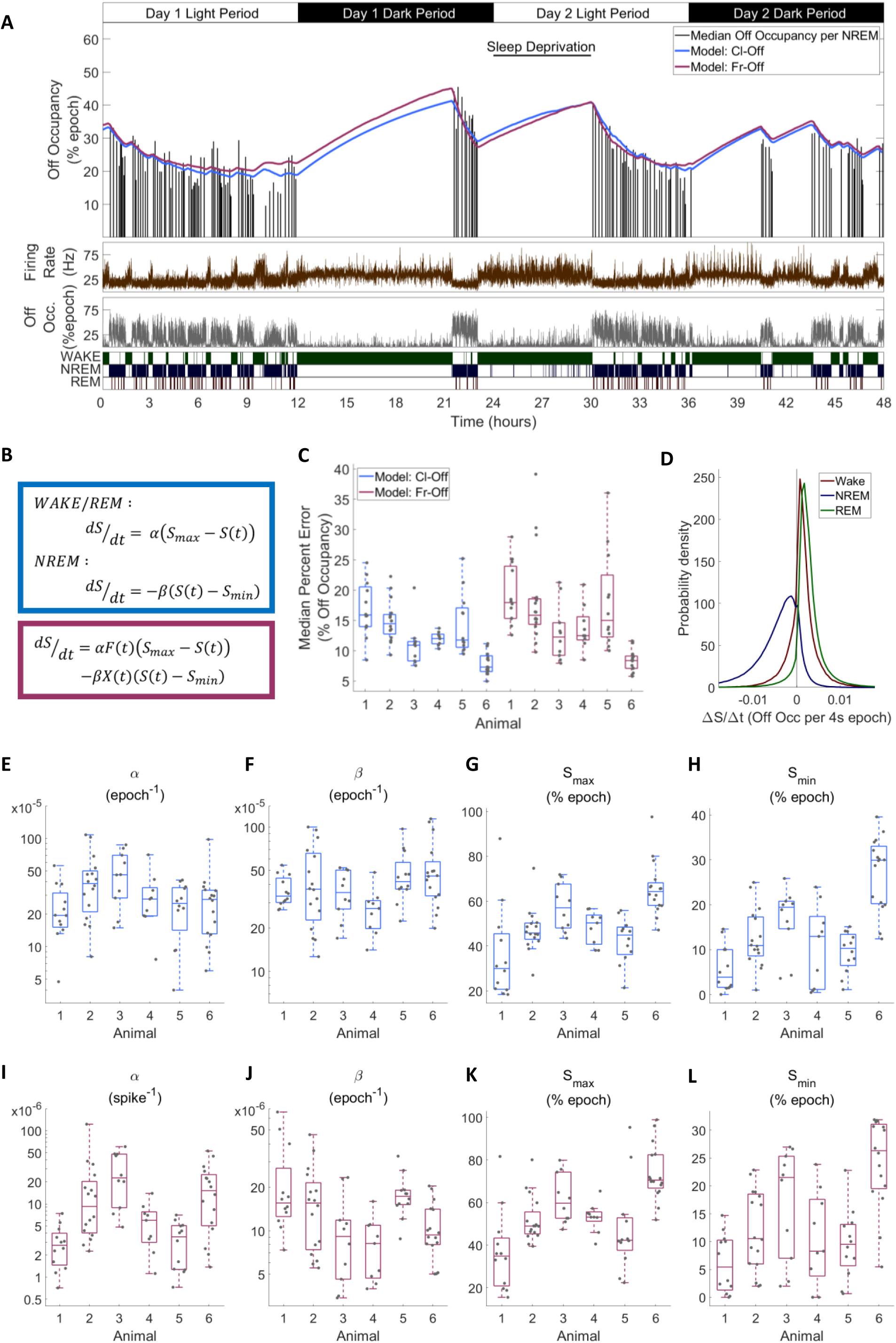
Process S is reflected in an LFP channel’s off occupancy and its dynamics are described well by both state-based and firing-rate-based models. **A)** An example of the novel model based on firing rates and off occupancy (purple), and the classic state-based model (blue), with optimised parameters describing the dynamics of off occupancy (median per NREM episode, black bars) over 48 hours. Sleep deprivation occurred as indicated at light onset of the second day and lasted 6 hours. Firing rate (brown), off occupancy (value per 4-s epoch, grey) and scored vigilance states are also shown. **B)** Equations for the classic state-based model (blue) and firing-rate-and-off-occupancy-based model for off occupancy (purple). **C)** For each animal, the distribution over channels of the median difference between the model and empirical off occupancy, expressed as a percentage of the off occupancy, for both classic and firing-rate-based models. **D)** The distribution of values of the change in Process S (***ΔS/Δt***) from one 4-s time step to the next derived from the Fr-SWA model in wake, NREM sleep and REM sleep. All mice, channels and time are pooled. **E-H)** The distribution of optimised values used for each parameter in the classic model and **I-L)** the firing rate model, with boxplots plotted separately for each animal. Source data is provided for this figure which contains all simulated Process S time courses, model parameters and the data required to reproduce them.

In this model, one term drives ***S*** towards an upper asymptote (***S_max_***) in proportion to firing rate (***F***), while the other drives ***S*** towards a lower asymptote (***S_min_***) in proportion to the off occupancy (***X***). Again, two rate parameters ***α*** & ***β*** are required. This model behaves similarly to previous models because firing is high in wake and low in NREM sleep, whereas off occupancy is high in NREM sleep and low in wake. This variant of the model is abbreviated as Fr-Off. Figure 5A also includes the fit from this model, demonstrating a high level of agreement between the two models and an accurate fit to the data.

The distribution of the median percent errors for the fits from both models over all animals and channels is shown in Figure 5C. As before, the model type and animal has a significant effect (Model: F_(1,138)_ = 8.06, p = 5.2×10^−3^; Animal: F_(5,138)_ = 17.2, p = 3.2×10^−13^; Model x Animal: F_(5,138)_ = 0.52, p = 0.76; two-way ANOVA with unequal groups) and the classic model achieved a slightly lower median percent error. However, this effect was again very weak compared with the variation in fit quality between animals and channels (Model: η^2^ = 0.034; Animal: η^2^ = 0.367; Model x Animal: η^2^ = 0.011; Channel (residuals): η^2^ = 0.588). Figure 5D shows the distribution of values for the change in modelled Process S from one simulated time step to the next resulting from the firing-rate-based model (Fr-Off), in wake, NREM and REM sleep, pooling all animals, channels and time. Unlike in the previous firing-rate-based model, REM sleep is now typically associated with an increase in Process S. The distributions of final optimised parameters are shown in Figure 5E-H for the classic model and Figure 5I-L for the firing rate and off occupancy model. Most parameters in both models were different between animals to a high significance level (Cl-Off ***α***: F_(5,74)_ = 2.4, p = 0.047; Cl-Off ***β***: F_(5,74)_ = 2.29, p = 0.055; Cl-Off ***S_max_***: F_(5,74)_ = 9.49, p = 6.4×10^−7^; Cl-Off ***S_min_***: F_(5,74)_ = 16.79, p = 8.0×10^−11^; Fr-Off ***α***: F_(5,74)_ = 9.86, p = 3.9×10^−7^; Fr-Off ***β***: F_(5,74)_ = 5.14, p = 4.6×10^−4^; Fr-Off ***S_max_***: F_(5,74)_ = 10.08, p = 2.8×10^−7^; Fr-Off ***S_min_***: F_(5,74)_ = 10.25, p = 2.3×10^−7^; one way ANOVA). Notably, the weakest evidence for inter-animal differences were for ***α*** and ***β*** in the classic model.

### Modelling identifies local variation in Process S dynamics

All four modelling approaches described here suggest the existence of variability in Process S between recording channels, indicating that a local component determines its dynamics. Figure 6A-D illustrates this diversity, showing the overlaid time courses of Process S for all channels within a single representative animal, expressed as a percentage of their individual maximum value for normalisation purposes. Although the models all fit empirical SWA or off period occupancy with generally high accuracy, there are nonetheless differences between models in the precise shape of Process S. The average Process S over all channels was also calculated for each model and compared with the Process S derived from applying the classical model to EEG SWA (Figure 6E). This reveals that Process S derived from modelling LFP SWA more closely resembles Process S derived from EEG SWA than Process S derived from off occupancy. This suggests that global Process S calculated at a higher spatial scale might reflect an averaging across a manifold of local Processes S that exist on a finer spatial level.

**Figure 6.**
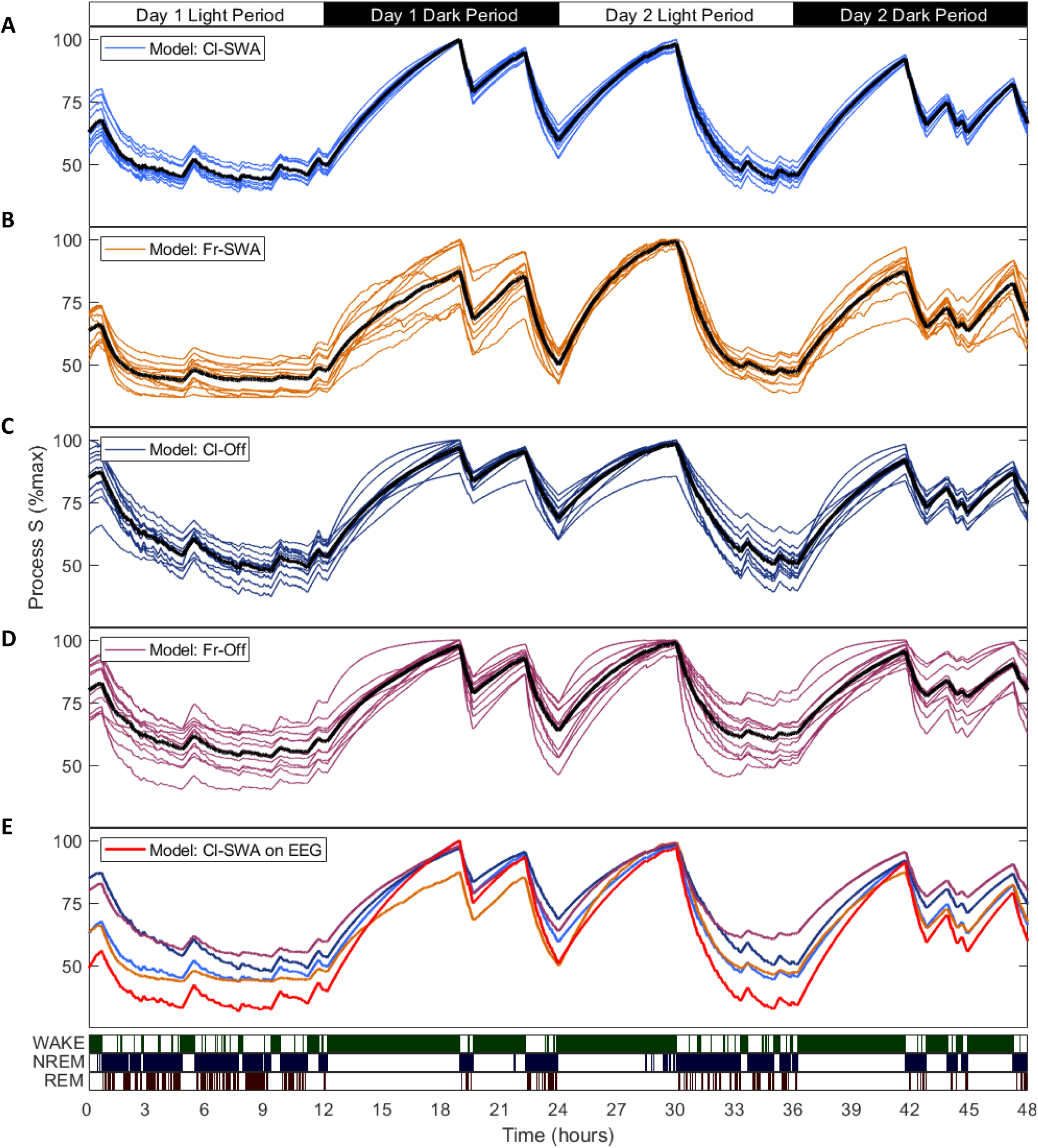
The time course of Process S is similar between models and individual channels. Process S time courses, expressed for each channel as a percentage of the maximum value, overlaid for all channels within a single representative animal obtained from **A)** the classic state-based model applied to LFP SWA, **B)** the firing-rate-based model applied to LFP SWA, **C)** the classic state-based model applied to off occupancy, **D)** the firing-rate-based model applied to off occupancy. In these panels the black line indicates the mean Process S over all channels. **E)** The mean Process S calculated over all channels is now plotted in colour (Cl-SWA light blue, Fr-SWA orange, Cl-Off dark blue, Fr-Off purple), and the red line indicates the Process S obtained by applying model Cl-SWA to the EEG derived SWA.

## Discussion

Here, we demonstrated that Process S dynamics can be derived entirely from neuronal spiking activity, without reference to global sleep-wake states. A model was outlined whereby the integrated history of local multi-unit firing rates, relative to a locally-defined firing rate set point, predicted the temporal dynamics of LFP and global EEG SWA. The accuracy of this model was demonstrated in a dataset of recordings from mouse frontal cortex over 48 hours, including both voluntary sleep and wake, and sleep deprivation. A novel metric for Process S was then presented, termed off occupancy, which measures the fraction of time a neural population spends in off periods, defined by the coincidence of LFP slow waves and multi-unit spiking silence. The modelling approach was combined with the off occupancy metric and tested on the same dataset to present a quantitative framework for understanding dynamics of sleep pressure at a highly local level in terms of neural spiking and off periods.

Central to this modelling perspective is the assumption that the generation of spikes by a neural network is in some way correlated with an increase in sleep pressure, and that this manifests in the subsequent expression of slow waves and off periods, reflecting Process S. The energetic cost of spiking is high (Attwell & Laughlin 2001), the regulation of a neurone’s firing rate set point is linked to cellular energetics (Styr et al., 2019; Vergara et al., 2019), and neurones are susceptible to cellular stresses that can result from sustained metabolic load, such as oxidative stress (Wang & Michaelis 2010; Cobley et al., 2018; Kempf et al., 2019). Furthermore, spiking activity may be mechanistically associated with synaptic plasticity, and it was suggested that firing rates are an important determinant of overall changes in synaptic strength, with higher spiking leading to greater changes (Graupner et al., 2016; Lappalainen et al., 2019). Compensatory processes exist within neurones to oppose cellular stress and related homeostatic challenges (Kültz 2005) and off periods could provide the opportunity for neurones to prioritise such processes, therefore mediating the restorative benefits of sleep (Vyazovskiy & Harris, 2013). Similarly, off periods are associated with distinct synaptic plasticity rules (Gonzalez-Rueda et al., 2018), and so their prevalence and patterning is likely also related to whatever regulation of synaptic strength occurs during sleep (Tononi & Cirelli 2014; Timofeev & Chauvette 2017; Seibt & Frank, 2019).

According to this view of sleep homeostasis, in principle, regulation can occur entirely at the local level, within cortical networks or perhaps even within single neurones. Indeed, this is reflected in our results through the differences between channels both with respect to the accuracy of the model and in the values of optimised parameters. Why then is global sleep preferred over a hypothetical state including asynchronous local off periods, which presumably could be used to sustain longer periods of behavioural wakefulness? It has been argued previously that it is ecologically optimal to synchronise off periods and undergo dedicated periods of total behavioural shutdown, because local off periods during waking impair behaviour (Vyazovskiy et al., 2011; Rattenborg et al., 2012). Mechanistically, the occurrence of off periods might be obstructed by strong synaptic coupling and shared neuromodulatory tone. Indeed, it has been observed that the degree of coupling between an individual neurone’s firing and the population firing rate, is variable between cells but characteristic to an individual neurone and likely reflects total synaptic strength with its neighbours (Okun et al., 2015). Some neurones may therefore be less able to express asynchronous off periods than others.

The preference for global sleep, despite its fundamentally local mechanisms, may be evidence that sleep homeostasis ultimately does not serve a single specific local function. Instead, the recent history of neuronal activity levels may regulate the local tendency to generate off periods (Process S), a signal which is integrated over larger neuronal populations through intrinsic network mechanisms in order to produce a global sleep propensity signal that estimates the total time spent awake with great accuracy. This mechanism would enable the brain to enforce a daily quota of sleep, which could have many benefits at the physiological and ecological level, rather than to initiate sleep in response to the homeostatic need of one specific regulated variable (Vyazovskiy, 2015). Importantly, in this mechanism, the variables which change as a function of time spent awake or asleep do not themselves need to be directly regulated by sleep. Conversely, variables which remain stable during continuous wake or sleep could still make essential contributions to the time-keeping mechanism. Note that this property is present in our model, since the levels of neural activity themselves are not directly proportional to the levels of Process S. The approximately equivalent overall accuracy of both firing-rate-based and vigilance-state-based models supports this possibility, as does the evidence that homeostatically regulated cellular variables can actually be stable during extended wakefulness, and that maintenance processes in sleep may be ultimately prophylactic (Vyazovskiy & Harris, 2013). In order to maximise the efficiency of global sleep, single neuronal activities may be modulated such that sleep pressure accumulates, on average, as uniformly as possible. This is consistent with recent reports that sleep regulates the population level firing rate distribution (Watson et al., 2016; Levenstein et al., 2017; Miyawaki & Diba, 2019) and could account for sleep’s link with synaptic and firing rate homeostasis, reconciling the local origins of sleep pressure accumulation with the global level of its dissipation.

Here, spike firing rate is used to represent the level of neural activity because it is convenient to record, locally variable, and directly linked to neuronal functionality. However, no strong claim is made that firing rates are necessarily causally responsible for the accumulation of sleep pressure. Indeed, it would likely be possible to obtain a reasonable quantitative account of sleep homeostasis using any physiological variable (or set of variables) that are consistently higher in either the wake or sleep state. For example, a model assuming that Process S increases in proportion to local cortical temperature, which, in laboratory rodents, drops by ∼2°C when falling asleep (Franken et al., 1991b; Tobler et al., 1997) and which has been mechanistically implicated in sleep regulation (Hoekstra et al., 2019), might also provide a plausible description of Process S dynamics. However, our results demonstrate that firing rates are a useful measurable correlate of the processes that directly underpin Process S, and therefore firing rate variance resulting from differences in experience and behaviour may well account for the variance in Process S accumulation in normal individuals, between waking periods and between cortical regions.

It should be noted that the firing-rate-based models used here typically slightly under-perform, relative to the classic model, in terms of minimising the error between simulation and empirical data. There are a number of reasons why a limit on the accuracy of the model is to be expected, related to technical restrictions rather than conceptual ones. In this approach, each MUA channel records only a few randomly sampled nearby neurones, which may have very different spiking properties, and so grouping these as a single measure of local network activity is somewhat arbitrary. It may be valuable to explore genetic and pharmacological manipulations of firing rate in order to further test this model’s validity. Unfortunately, the effects of any manipulation on the accumulation, expression and dissipation of sleep pressure, separately to the effects on firing, is unknown, and so this may be hard to interpret. For example, a recent study found that systemic atropine administered during behavioural wakefulness produces slow wave activity, reducing spiking (as measured by c-Fos) yet increasing the duration of subsequent NREM sleep (Qiu et al., 2015). While this was interpreted as evidence that spiking activity is not related to sleep pressure accumulation, the direct effects of atropine on Process S and the functional significance of the resulting induced slow oscillation are unclear. Similarly, another study found that local optogenetic activation of cortex during sleep, despite raising firing rates to waking levels, was not accompanied with increased SWA or off periods during subsequent NREM sleep (Rodriguez et al., 2016). Again, it is difficult to disentangle the direct effects of this local stimulation, which is not physiologically realistic, on the mechanisms surrounding the accumulation, expression and dissipation of sleep pressure. On the other hand, these results might be interpreted as evidence that the expression of the level of Process S involves an integration of homeostatic sleep need across neural populations and therefore local firing rate manipulations are not able to substantially influence Process S dynamics at a global level.

It is reasonable to assume that sleep homeostasis unfolds on multiple time scales and Process S as defined by these models describes a relatively fast one, approaching its upper asymptote after continuous wakefulness on a time scale of hours. The inclusion of processes acting over longer or shorter time scales might explain discrepancies in all these models, however, the challenge remains to identify what these could be. Furthermore, the role of REM sleep in Process S dynamics has not been explicitly addressed or considered in the construction of these models. Depending on the model variant, REM sleep is associated either with a small increase or small decrease in Process S, because firing rates are low in REM sleep (closer to NREM sleep than waking) and yet off period occupancy is also low (closer to wake levels than NREM sleep). It is possible that REM sleep might represent a homeostatically neutral state, in which the level of Process S changes minimally, or not at all (Vyazovskiy & Delogu, 2014).

In summary, our data suggest that Process S is reflected at multiple scales in the brain, from EEG and LFP slow wave activity to the occurrence of local multi-unit off periods. Its dynamics across all scales can be described quantitatively using information derived only from local neuronal activity with comparable accuracy to the classical model of Process S which depends on global sleep-wake history. We postulate that, despite its local origins, Process S may be integrated across networks for the purpose of tracking time spent awake and used to enforce a daily quota of global sleep, rather than serving the regulation of specific physiological variables.

## Funding acknowledgements

BBSRC (BB/K011847/1, BB/M011224/1), MRC (MR/L003635/1), Wellcome Trust (098461/Z/12/Z), John Fell OUP Research Fund (131/032).

